# TIF1γ regulates stability of T regulatory cells during inflammation

**DOI:** 10.1101/2024.05.23.595623

**Authors:** Eugenio Contreras-Castillo, Jesús Daniel Zambrano-Romero, Diego Pérez-Vázquez, María Guadalupe García-Patiño, Gustavo Tapia-Urzúa, Aurora Candelario-Martínez, Jose Luis Ramos-Balderas, Irma Ileana Licona-Limón, Iris Madera-Salcedo, Florencia Rosetti-Scuitto, Jose Carlos Crispin-Acuña, Lilia Noriega-López, Porfirio Nava, Félix Recillas-Targa, Veesa Kaartinen, Jorge Henao-Mejía, Richard Flavell, Paula Licona-Limón

## Abstract

T regulatory cells are a suppressor population critical to control inflammation and maintain tissue homeostasis. The TGF-β pathway is a fundamental signal for T regulatory cell differentiation, yet the molecular determinants of how TGF-β signals regulate all these processes are not completely understood. TIF1γ was demonstrated to promote a noncanonical TGF-β/Smad pathway in HSC, however the role of TIF1γ in Treg function has been largely ignored. Here we showed that TIF1γ deficient Tregs lose stability and acquire an effector phenotype in the presence of an inflammatory stimuli or upon activation. TIF1γ deficient T regs gain a Th1-like Treg phenotype or become pro-inflammatory exTregs, by losing Foxp3 expression and acquiring IFNγ expression in an autoimmune model. Loss of TIF1γ in Tregs was cell-intrinsic and was also accompanied by increased proliferation and enhanced glycolytic capacity upon activation. Additionally, we demonstrated that in the absence of TIF1γ, there was an increased methylation status in the CNS2 enhancer region of the Foxp3 locus, further suggesting increased susceptibility for loss of Foxp3 expression. Mechanistically, TGF-β was in part responsible to inhibit the Th1-like bias a Treg has in the absence of TIF1γ, however, the proliferative phenotype observed in these cells was mostly independent of TGF-β signaling. Finally, we identified the beta-catenin pathway as the molecular mechanism driving both TIF1γ-dependent Treg stability and proliferation upon inflammation. Altogether, our data demonstrated that TIF1γ is required for the maintenance of a suppressor phenotype and stability of Treg lymphocytes during inflammatory conditions *in vivo* and represents a new modulatory pathway to manipulate Treg cells for therapeutic purposes.

---

TIF1γ or Trim 33 (Transcriptional intermediary factor 1 γ or Tripartite motif containing 33) is a ubiquitin ligase belonging to the Trim protein family which was recently shown to promote a noncanonical response to TGF-β in HSC (1). TGF-β is a critical cytokine for T reg cell differentiation and function (2–8), the TGF-β canonical signaling pathway depends on the phosphorylation of Smad2 and Smad3, which interacts with Smad4 to regulate gene expression. The identification of TIF1γ as an interacting partner for Smad2/3, demonstrated an alternative pathway in which Smad2/3/TIF1γ complexes could lead to a different gene expression profile and cell fate, compared to the Smad4-dependent pathway in HSCs (9–11). The regulation of Treg cells through the canonical TGF-β pathway has been extensively explored (12–15), however, the role of TIF1γ in Tregs has been largely ignored (16) hence our objective was to characterize the role of the TIF1γ pathway in the differentiation, stability and function of Tregs. Using different conditional deficient mouse models (cKO), we demonstrated that under homeostatic conditions, TIF1γ is dispensable for Treg generation; however, these cells showed defective suppressor capacities *ex-vivo* as well as differential expression of some Treg functional markers. Moreover, aged TIF1γ cKO mice develop an increased inflammatory phenotype, with decreased body weight, decreased colon length, and increased INFγ expression in pTregs. Interestingly, upon inflammation, our data demonstrated that TIF1γ is required for the maintenance of suppressive functions and stability of Tregs, both *in vivo* and *in vitro.* In a T cell transfer colitis model, TIF1γ deficient Tregs lose expression of Foxp3 and display a Th1-like phenotype characterized by Tbet and IFNγ expression. We found that TIF1γ deficient Tregs acquire a glycolytic metabolic profile and develop increased proliferative capacity; further promoting wasting syndrome and inflammation in the colon. Moreover, we demonstrated that TIF1γ is necessary to prevent the acquisition of a Th1-like phenotype in Tregs in an autoimmune model. Additionally, we identified increased expression and activation of beta catenin in TIF1γ-deficient Treg cells, most likely due to decreased proteosomal degradation promoted by TIF1γ (17), which was also in part responsible for promoting loss of Foxp3 and proliferation in Tregs. Impaired stability of Tregs lacking TIF1γ was also confirmed by an increased methylation status at the CNS2 enhancer region of the Foxp3 locus. Our study provides strong evidence that TIF1γ is required for the maintenance of a suppressor phenotype and stability of Treg cells upon inflammation and centers TIF1γ as an interesting target to regulate Treg cell plasticity and commitment *in vivo*.

## Results

### T cell-specific TIF1γ aged deficient mice showed signs of increased inflammation and increased Th1-Like Tregs

To evaluate the role of TIF1γ in Tregs we generated a conditional deficient TIF1γ mouse model (TIF1γ cKO) which was also a reporter for Foxp3, by crossing a TIF1γ floxed mice (18) with a CD4 Cre line (19), similar to what has been recently reported (16), but in the Foxp3 GFP background (20) to delete TIF1γ in total T cells and to be able to track Treg cells *in vivo*. At steady state, these mice were viable and fertile and showed no major phenotypic defects in T cell development, measured as frequencies of CD4, CD8 and CD4+ Foxp3+ Treg (Extended data Fig 1a) in 8 weeks old mice; however, Tregs from TIF1γ cKO young mice showed lower levels of CD73 and higher levels of CD25, both important functional markers of Tregs (Extended data Figure 1b). Considering Tregs are central effectors in restraining age-related inflammation (21), we also evaluated the phenotype of TIF1γ cKO aged mice (>40 weeks). We observed signs of increased inflammation such as shortened colon length, decreased body weight, decreased frequencies of Foxp3^+^ cells, and increased frequencies of resting Tregs (CD44^low^CD62L^+^) (22) (Fig 1a-d), suggesting a possible defective activation of Tregs in the absence of TIF1γ in aged mice. Interestingly, Tregs from aged TIF1γ cKO mice showed a higher expression of IFNγ within the Th1-Like Treg population defined as Foxp3^+^IFNγ^+^ (Fig 1e), although the frequencies of these cells were similar between mice groups (Extended data 1c). These cells are typically only present in Type 1 infectious/inflammatory models (23), expression of IFNγ in Tregs has been associated with a better suppressor capacity towards Type 1 inflammation (24) but has also been associated with a decreased suppressor function (25), hence we asked whether Tregs from TIF1γ cKO mice could have a different suppressor capacity.

**Fig 1.**
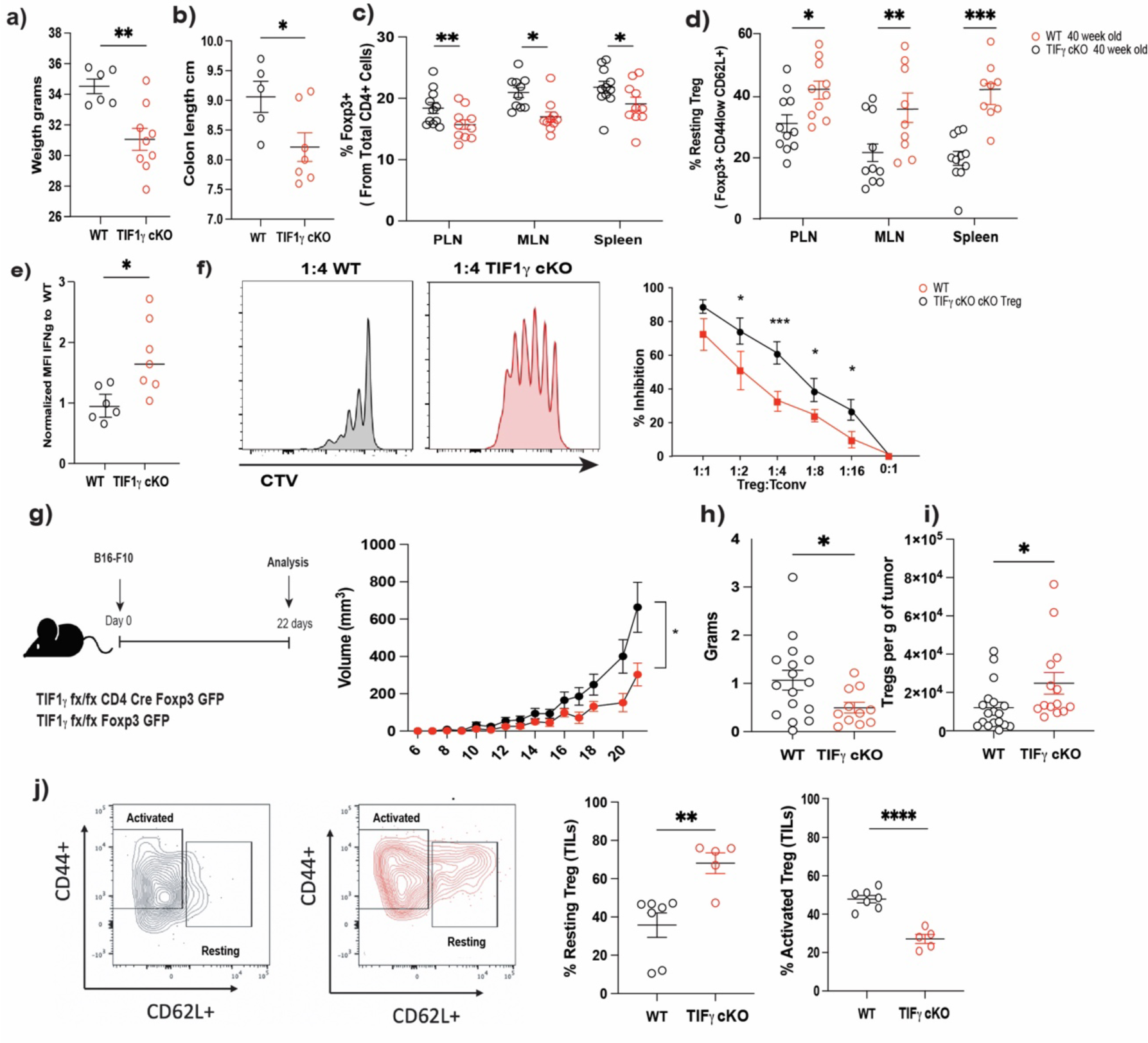
TIF1γ is required for a proper Treg suppressor capacity and activation *in vivo*. a)Body weight at 40 weeks of age in TIF1γ WT and cKO mice. b) Colon length in 40 weeks old WT and TIF1γ cKO mice. c) Frequency of CD4^+^Foxp3^+^ cells in peripheral lymph node (PLN), mesenteric lymph node (MLN) and spleen of 40 week old WT and TIF1γ cKO mice. d) Frequency of resting Tregs (CD4^+^Foxp3^+^CD62L^+^CD44^-^) in PLN, MLN and spleen of 40 week old WT and TIF1γ cKO mice. e) IFNγ Mean Fluorescence Intensity in TIF1γ cKO splenic Tregs normalized to WT Tregs from 40 week old mice. f) Representative histogram at 1:4 ratio co-cultures. Percentage of inhibition *in vitro* suppression assay of WT and TIF1γ cKO Tregs co-cultured with conventional T cells at different ratios. g) Experimental design of the B16 melanoma model and tumor growth in WT and TIF1γ cKO mice. h) Tumor mass in WT and TIF1γ cKO mice at day 22 post B16 injection. i) Number of Foxp3+ cells in the tumor infiltrating lymphocytes (TIL). j) Representative contour plot and frequencies of resting (CD4^+^Foxp3^+^CD62L^+^CD44^-^) and activated Tregs (CD4^+^Foxp3^+^CD62L^-^ CD44^+^) in tumor infiltrating lymphocytes in WT and TIF1γ cKO mice at day 22 post B16 injection. n= 6-16 mice per group from at least three independent experiments. p values * p<0.05, ** p<0.01 and *** p<0.001, **** p<0.0001

### TIF1γ deficiency impairs pTregs suppressor capacity and iTreg differentiation *in vitro*

To address the functional capacity of Tregs *ex vivo*, we performed *in vitro* suppression assays by co-culturing responder naive T cells with pTregs at different ratios, our data demonstrated that in the absence of TIF1γ, Tregs have a decreased suppressor capacity with higher frequencies of proliferating responder T cells in these co-cultures in most of the ratios analyzed (Fig 1f). We next asked if the absence of TIF1γ was affecting the potential for a naive cell to become iTreg. To explore this hypothesis, we isolated naive T cells from WT or TIF1γ cKO mice and cultured these cells in polarizing conditions towards Treg differentiation. Our data showed that lower frequencies and numbers of iTregs (CD25^+^Foxp3^+^) were generated from TIF1γ deficient naive T cells (Extended data Figure 1d) and consequently TIF1γ is required for an optimal iTreg differentiation *in vitro*. We also observed an increased frequency of Th1-like Tregs in cultures from TIF1γ cKO cells, with increased expression of T-bet and IFNγ compared to WT iTregs (Extended data 1e). To explore possible reasons for decreased iTreg differentiation in cultures from TIF1γ cKO mice, we analyzed proliferation and apoptosis in these cultures. Interestingly TIF1γ deficient iTreg cultures had an increased proliferative capacity but no difference was observed in apoptosis induction (extended data Fig 1f, 1g), hence the decreased frequency of iTregs in TIF1γ cKO cultures is not related to decreased proliferation or increased death in the

Treg population. It is important to note that it seems that the phenotype observed in TIF1γ cKO Tregs, requires either activation of the cells or an inflammatory stimulus (phenotype in old mice) to take place, suggesting that TIF1γ might be required for sustained stability of Tregs in a context-dependent manner. To validate this hypothesis, we isolated pTregs from WT and cKO mice and cultured these cells for 5 days in the presence of anti-CD3, anti-CD28 and IL-2 plus different inflammatory cytokines (IL-4, IL-6, IFNγ or TNFα), to evaluate the frequency of Foxp3 expressing cells in these conditions. Our data clearly demonstrated, that upon activation and in the presence of IL-2, IFNγ and TNFα, TIF1γ cKO Treg express significantly lower levels of Foxp3 compared to their WT counterparts (extended data 1h), we also observed increased proliferative capacity of cKO Tregs in these conditions (extended data 1i), consequently upon activation and in the presence of inflammatory stimulus, TIF1γ is required for a stable Foxp3 expression and decreased proliferative capacity of Tregs.

### TIF1γ cKO mice showed increased anti-tumor response

Considering the decreased suppressor capacity in TIF1γ cKO Tregs observed in the *in vitro* suppression assays, we then evaluated if, in an *in vivo* model, we could also observe a diminished suppressor function in these cells. We challenged WT and TIF1γ cKO mice with the B16 mouse melanoma cell line, followed the tumor growth, and analyzed the Treg populations in draining lymph nodes (DLN), non-draining lymph nodes (NDLN), spleen, and tumor-infiltrating lymphocytes (TILs). A reduction in tumor growth was observed in TIF1γ cKO compared to WT mice (Fig 1g) as well as a decreased tumor mass (Fig 1h). When we evaluated the frequencies and absolute numbers of Tregs in this model, we only found a significant difference in the numbers of Tregs within the TILS (Fig 1i), however, there was no difference in the ratio between CD8s and Tregs (extended data 1j). Additionally, we did not find any difference in some of the functional markers previously evaluated in pTregs from 8-week-old mice at steady state, when we evaluated the TILs (Extended data 1k) yet we did observe an increased resting phenotype on TIF1γ cKO Tregs within the tumor (Fig 1j). These data suggest a possible defective suppressor capacity in TIF1γ cKO Tregs which could lead to a better anti-tumoral response; however, considering that these experiments were performed in TIF1γ cKO with the CD4-Cre transgene, where all T cells (including CD4 helper and CD8 cytotoxic T cells) lack the expression of TIF1γ, we cannot exclude the contribution of these other cells in addition to Tregs in the observed phenotype, hence we decided to explore the suppressor capacity of Tregs in a cleaner experimental setting.

### TIF1γ deficient Tregs lose Foxp3 and become Th1-like during an autoimmune response

We next sought to evaluate the phenotype of TIF1γ cKO Tregs in an adoptive transfer model of colitis (26). We isolated congenically marked CD45.1 naive T cells from WT mice and single transferred or co-transferred these cells with CD45.2 WT or TIF1γ cKO Tregs into Rag2 deficient hosts. 6-8 weeks post-transfer, mice receiving TIF1γ cKO Tregs exhibited increased weight loss and increased frequencies of transferred cells expressing IFNγ+, similar to the single transferred naive group (Fig 2a-b), supporting the notion of a deficient suppressor capacity in TIF1γ deficient Tregs. We also observed an increased clinical score in the colon with loss of intestinal structure in the co-transfered group with TIF1γ cKO Tregs that was similar to the single transfer naive group (Fig 2c). We next evaluated the phenotype of transferred Tregs in this model, as expected, control Tregs prevented weight loss and colon damage (Fig 2a and 2c) and maintained high frequencies of Foxp3 with only a few cells losing this marker upon transfer, whereas TIF1γ cKO Tregs showed a decreased frequency of Foxp3 expressing cells and an increased frequency of ex-Tregs (CD45.2^+^Foxp3^-^) in the spleen and MLN of Rag ^-/-^ hosts (Fig 2d). Expression level of Foxp3 within the Foxp3+ population was not different between groups (Extended data 2a), however, the acquisition of IFNγ expression was significantly increased in TIF1γ deficient Tregs compared to WT Tregs (Fig 2e). Interestingly, when we evaluated the expression of IFNγ on the ex-Treg population (cells losing Foxp3 expression) in these mice, we found that compared to WT Tregs, TIF1γ cKO tregs showed increased frequencies of IFNγ^+^ cells (WT 5.01+/-1.42 vs 14.31+/-3.54), acquiring a Th1 phenotype *in vivo* (Fig 2f). These data suggest that TIF1γ deficient Tregs had decreased stability and became Th1-like or even lost Foxp3 expression upon exposure to inflammatory signals *in vivo*.

**Fig 2.**
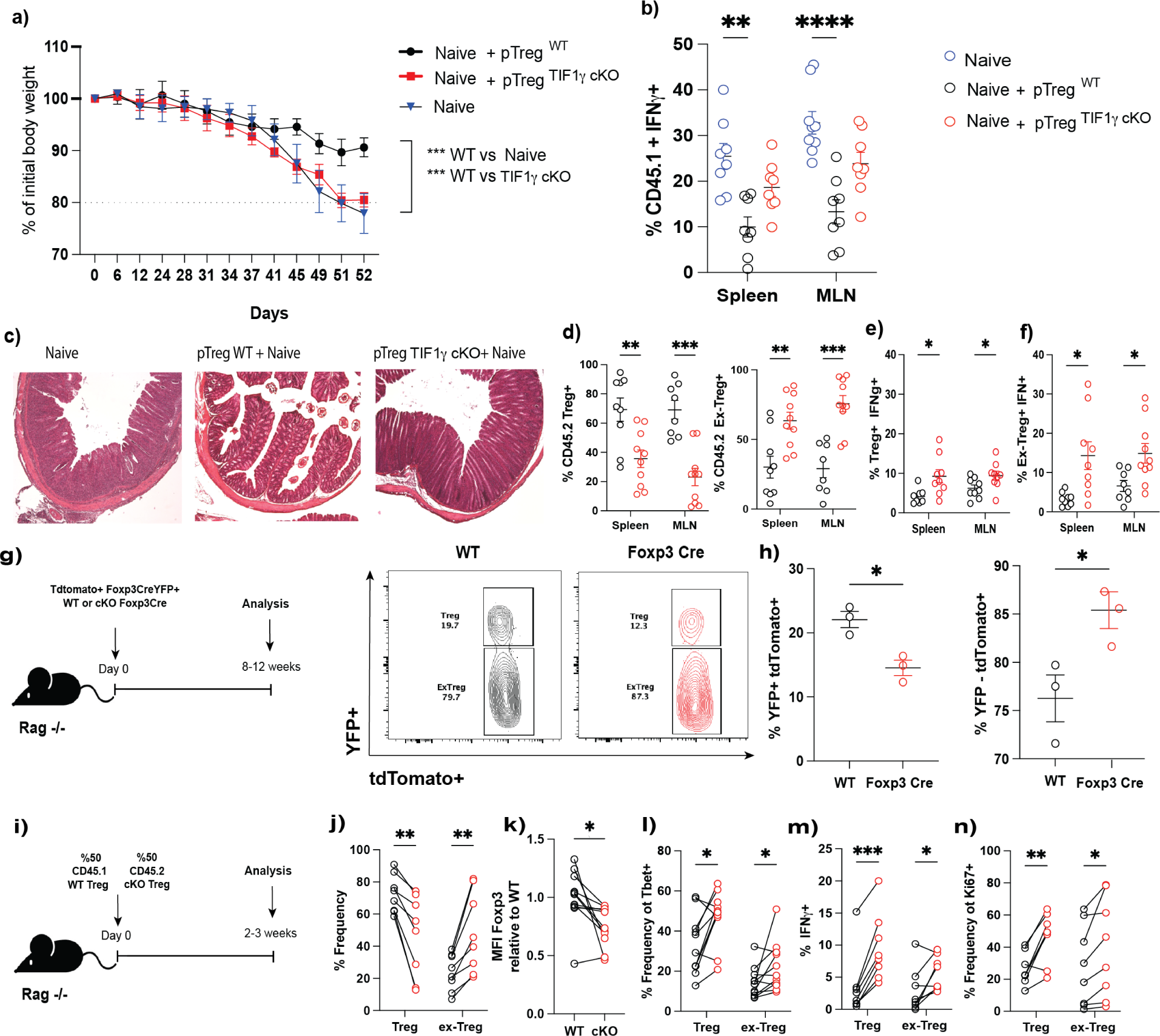
TIF1γ deficient Tregs lose Foxp3 expression and acquire a Th1-like phenotype upon transfer into Rag-/- mice. a) Body weight loss in Rag^-/-^ single transferred or co-transfered with conventional naive T cells (CD4^+^CD25^-^CD62L^+^CD44^-^CD45.1^+^) and WT or TIF1γ cKO Tregs (CD4^+^CD25^+^GFP^+^CD45.2^+^). b) Frequency of conventional naive T cells expressing IFNγ in Rag^-/-^ transferred mice. c) Colon histology of transferred Rag -/- mice. d) Frequencies of WT or TIF1γcKO Tregs (CD4^+^Foxp3^+^CD45.2^+^) and ex-Tregs (CD4^+^Foxp3^-^CD45.2^+^) in Rag^-/-^ transferred mice. e) Frequencies of WT or TIF1γ cKO Tregs expressing IFNγ in Rag^-/-^ transferred mice. f) Frequencies of WT or TIF1γ cKO ex-Tregs expressing IFNγ in Rag^-/-^ transferred mice. g) Experimental design of single transferred WT or TIF1γ cKO fate Tregs into Rag^-/-^ hosts, representative density plot of Tregs (CD4^+^Foxp3^-^YFP^+^Tomato^+^) and exTregs (CD4^+^Foxp3^-^YFP^-^ Tomato^+^) at 8-week post-transfer. h) Frequencies of WT or TIF1γ cKO Tregs (YFP^+^tdTomato^+^) and ex-Tregs (YFP^-^tdTomato^+^) in the spleen of Rag^-/-^ transferred mice at week 8 post-transfer. i) Experimental design of co-transferred WT (CD4^+^CD25^+^GFP^+^CD45.1^+^) and TIF1γ cKO (CD4^+^CD25^+^GFP^+^CD45.2^+^) Tregs into Rag^-/-^ hosts. j) Frequencies of WT or TIF1γ cKO Tregs and exTregs in the spleen at 3 weeks post-transfer into Rag^-/-^ hosts. k) Foxp3 mean fluorescence intensity of WT or TIF1γ cKO Tregs in the spleen at 3 weeks post-transfer into Rag^-/-^ hosts. l) Frequencies of WT or TIF1γ cKO Tregs and exTregs expressing Tbet in Rag^-/-^ transferred mice. m) Frequencies of WT or TIF1γ cKO Tregs and ex-Tregs expressing IFNγ in Rag^-/-^ transferred mice. n) Frequencies of WT or TIF1γ cKO Tregs and ex-Tregs proliferating in Rag^-/-^ transferred mice. n=10-11, (except g) were n=3 per phenotype and condition). Results from at least two-three independent experiments. * p<0.05, ** p<0.01, *** p<0.001, **** p<0.0001.

### Treg stability loss in the absence of TIF1γ is cell intrinsic

To determine if the phenotype we observed on Tregs was cell intrinsic, we fate-mapped single transferred WT or TIF1γ cKO fate reporter Tregs, isolated from a Rosa26 td-Tomato TIF1γ wt/wt Foxp3 Cre YFP or a Rosa26 td-Tomato TIF1γ flox/flox Foxp3 Cre YFP mice, respectively (27–28), into Rag ^-/-^ hosts (Fig 2g). 8-12 weeks post-transfer we analyze absolute numbers and frequencies of transferred cells in the spleen, we also follow the weight loss over the course of the experiment. Our data showed no significant difference in weight loss upon single Treg transfer independently of the phenotype of these cells (Extended data 2b), however, we did observe an increased expansion of TIF1γ deficient Tregs compared to WT Tregs, both within the Tregs and ex-Tregs populations (Extended data 2c), similar to what we observed in our *in vitro* iTreg cultures (Extended data 1f). Additionally, TIF1γ deficient Tregs showed increased frequencies of ex-Tregs (Rosa td-Tomato^+^YFP^-^) and decreased frequency of Tregs (Rosa td-Tomato^+^YFP^+^) when compared to single transferred WT fate reporter Tregs cells (Fig 2h). Collectively these data suggest that the loss of Foxp3 expression in the absence of TIF1γ is cell intrinsic and does not require an inflammatory microenvironment to occur. We then wanted to assess the impact of TIF1γ deficiency in the T cell capacity to maintain a Treg phenotype once differentiated *in vivo*, to do this we performed single transfer experiments of naive WT or TIF1γ cKO fate reporter T cells into Rag -/- recipients and followed the generation and maintenance of Foxp3 expression in those cells (Extended data 2d). Treg cell (Rosa^+^YFP^+^) frequencies in spleen and MLN of mice transferred with TIF1γ cKO T cells showed a trend towards reduction compared to WT donor cells; while ex-Treg (Rosa^+^YFP^-^) cells showed a trend towards increase in the TIF1γ deficient transferred group (Extended data 2e). In sum, these data suggest that the lack of stability of TIF1γ cKO could occur *in vivo* even in d*e novo* generated pTregs cells and might not be restricted to the developmental origin of these cells, however further experiments are required to confirm this conclusion.

### TIF1γ cKO Tregs loss stability even in the presence of functional Tregs

To further evaluate if the phenotype observed on TIF1γ deficient Tregs can be controlled by bonafide Tregs *in vivo*, we isolated CD45.1 WT Tregs and CD45.2 TIF1γ deficient Tregs from WT or TIF1γ cKO fate mice and we co-transferred these cells into Rag -/- hosts (Figure 2i). 2-3 weeks post-transfer we evaluated the phenotype of Tregs in spleen and mesenteric lymph nodes and found a significantly decreased frequency of Foxp3^+^ cells in TIF1γ deficient Tregs compared to WT Tregs as well as increased frequencies of ex-Tregs (Foxp3^-^) cells (Fig 2j and Extended data 2f). We also observed a reduced Foxp3 expression in TIF1γ cKO Tregs compared to WT counterparts (2k). We then analyzed the capacity of these transferred cells to become Th1-like and found a significant increase in the frequencies of Tbet and IFNγ Tregs from TIF1γ deficient donors, as well as increased frequency of Tbet and IFNγ in Tregs that have already lost Foxp3+ (Th1 ex Tregs), in TIF1γ deficient Treg donor cells (Fig 2l-m). Interestingly, Treg and exTregs cells derived from TIF1γ cKO mice also showed increased proliferative capacity *in vivo* (Fig 2n). Decreased frequency and numbers of Tregs and increased frequency of exTregs, was also confirmed using Tregs from CD4 Cre TIF1γ flox/flox mice (Extended data 2g-h), demonstrating that the phenotype observed in Tregs, is independent of the Cre mouse line used in our transfers. All together these data support the notion that in the absence of TIF1γ, Tregs experience decreased stability and a higher susceptibility to acquire a Th1-like phenotype in a cell transfer model, even in the presence of functional Tregs.

### TIF1γ- deficient T regs cells have an increased methylation status on the CNS2 region

DNA demethylation on the CNS2 (conserved noncoding sequence 2) enhancer region of the *Foxp3* locus has been largely associated with a stable T regulatory phenotype (29); hence we asked whether the phenotype observed in Treg instability in the absence of TIF1γ, could also be related to the methylation status on this region. We isolated naive, pTregs and iTregs from WT and TIF1γ cKO and performed a CpG methylation analysis (22, 30). Our data demonstrate an increased methylation status on the CNS2 region of pTregs and iTregs, but not on the promoter region in cells isolated from TIF1γ cKO mice compared to WT Tregs (Extended data 3b-3g). These data further support the hypothesis that TIF1γ is important to maintain Treg stability and to prevent the acquisition of a Th1-like phenotype on Tregs upon inflammation. Accordingly, the stability hallmark of stable Tregs is also affected in the absence of TIF1γ, however the exact mechanism involved in this phenotype warrants further investigation.

### TGF-β controls part of the phenotype observed in TIF1γ cKO Tregs

We next sought to explore the molecular mechanism controlled by TIF1γ in Tregs, responsible for the maintenance of Treg stability and function. TIF1γ has been shown to mediate a non-canonical TGF-β/Smad pathway (9,10); however, it also promotes β-catenin degradation through its E3 ubiquitin ligase domain acting as an inhibitor of this pathway (17). Both the TGF-β and the β-catenin pathways have been involved in the regulation of functional responses in Tregs (25, 31–32), therefore we asked which one of these two possible pathways was responsible for the phenotype observed in TIF1γ deficient Tregs. We isolated pTregs from WT or TIF1γ cKO mice and we activated these cells for 5 days with anti-CD3, anti-CD28, and IL-2 in the presence of TGF-β or neutralizing antibodies against TGF-β, as well as in the presence of IFNγ as a way to mimic an inflammatory signal *in vitro*. Our results showed that the increased proliferative capacity of TIF1γ deficient Tregs was mostly independent of TGF-β (Fig 3a), suggesting that the anti-proliferative canonical effect of TGF-β in T cells seems to be dependent of TIF1γ. We then asked if the acquisition of a Th1-like phenotype in Tregs lacking TIF1γ was controlled by the TGF-β pathway, our data showed that such acquisition was inhibited when TGF-β was present while inhibition of TGF-β by using a neutralizing antibody, further enhanced the bias towards Th1 (Fig 3b). All together these data suggest that TGF-β and TIF1γ are directly linked to inhibit proliferation in Tregs, while the acquisition towards a Th1-like phenotype inhibited by TGF-β was TIF1γ- independent.

**Fig 3.**
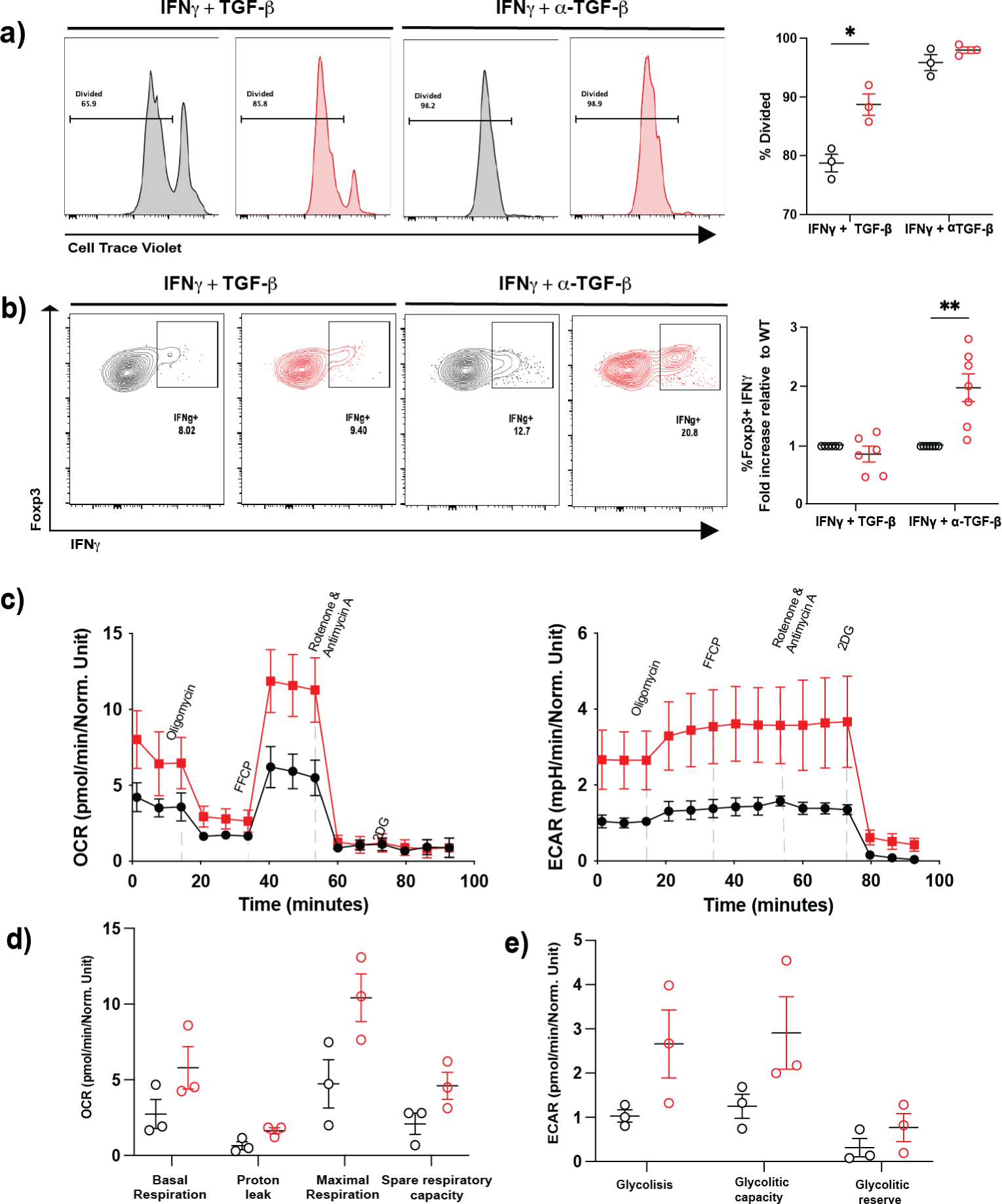
TIF1γ cKO Tregs have TGF-β-dependent and TGF-β-independent control of proliferation and stability, as well as an impaired metabolic profile. a) pTregs isolated from WT or TIF1γ cKO mice were stained with CTV and stimulated ex-vivo for 5 days with anti-CD3, anti-CD28, IL-2 and IFNγ, in the presence of TGF-β or a neutralizing antibody against TGF-β. Representative histogram of CTV dilution and frequencies of divided pTregs in culture. b) Representative contour plots with the frequencies of IFNγ and Foxp3 expressing Tregs in cultures described in a). Frequencies of Foxp3^+^IFNγ^+^ Tregs presented as fold increase relative to WT Tregs. c) Oxygen rate consumption (OCR) and extracellular acidification rate (ECAR) of WT or TIF1γ cKO iTregs measured in a Seahorse analyzer. d) Mean and standard error of basal respiration, proton leak, maximal respiration and spare respiratory capacity of WT and TIF1γ cKO iTregs. e) Mean and standard deviation of glycolysis, glycolytic capacity and glycolytic reserve of WT and TIF1γ cKO iTregs. n=3-9 mice per group from at least three independent experiments. p values * p<0.05 and ** p<0.01

### TIF1γ cKO iTregs acquire a glycolytic phenotype

A characteristic feature of a Treg cell versus any effector T cell is the metabolic profile they have (33), that is intimately related to the proliferative capacity they typically show, being effector T cells more glycolytic and Tregs (along with naive and memory T cells) more respiratory (oxidative phosphorylation). With this idea in mind, and considering the phenotype observed in TIF1γ deficient Tregs, we evaluated the metabolic profile of WT and TIF1γ cKO iTreg cells using a Seahorse analyzer (34). Interestingly, our data provided additional evidence regarding a higher proliferative capacity in Tregs lacking TIF1γ expression, by showing increased glycolytic activity when compared to WT Treg cells which showed mostly respiratory capacity (Fig 3c). TIF1γ cKO Tregs also showed an increased respiratory capacity compared to Wt Tregs (Fig 3c). Analysis of different parameters associated with the respiratory capacity in a cell was also increased on TIF1γ cKO Tregs compared with WT Tregs (Fig 3d), as well as those related with a higher glycolytic capacity (Fig 3e), hence the absence of TIF1γ, Tregs seems to improve their respiratory capacity but at the same time they enhance their glycolytic potential. Accordingly, TIF1γ cKO Tregs have a metabolic profile that fits with the proliferative demand observed in these cells. In sum, these data suggest that in the absence of TIF1γ, Tregs acquire a higher proliferative capacity that seems to be partially dependent of the TGF-β signaling pathway, and that these cells increase their respiratory capacity while developing a higher glycolytic potential, that closely resembles the metabolic profile observed in Th1 effector cells.

### TIF1γ regulates β-catenin abundance and promotes the stability of Tregs

Finally, we wanted to explore whether part of the phenotype in TIF1γ deficient Tregs was dependent on the regulation of the β-catenin pathway. To answer this question, we analyzed the expression of β-catenin in pTregs and iTregs from WT and TIF1γ cKO (Foxp3Cre fate) activated *in vitro*. We observed higher expression of β-catenin in TIF1γ deficient Tregs irrespective of their origin (Fig 4a), as well as a statistically significant increase in TCF7 (a downstream target of β-catenin) in these cells (Fig 4b). We further confirmed increased active beta-catenin (ABC) in TIF1γ cKO Tregs compared to Wt Tregs by WB (Fig 4c). We next wanted to investigate the influence of increased β-catenin levels in the phenotype observed on TIF1γ deficient Tregs and performed *in vitro* iTreg cultures in the presence or absence of XAV 939 (35) (a stabilizer of the β-catenin degradation complex: Axin:CK1a:GSK3b:APC). Our data demonstrated that the promotion of β-catenin degradation on TIF1γ cKO Tregs, was enough to ablate the increased frequency of Th1-like Tregs previously observed (Fig 4d) and it also decreased the enhanced proliferation observed in these cells (Fig 4e). To further demonstrate that the β-catenin pathway was indeed, at least one the mechanisms responsible for the lack of stability in TIF1γ cKO Treg cells *in vivo*, we co-transferred CD45.1 WT Tregs and CD45.2 TIF1γ deficient Tregs into Rag ^-/-^ mice and treated those animals with 10mg/kg IP of XAV 939 or vehicle every 4 days (Fig 4f). As expected, TIF1γ cKO Tregs lose Foxp3 and become exTregs in the control group (vehicle) (Fig 2j and Fig 4g). In contrast, the addition of XAV 939 completely abrogated the loss of Foxp3 in TIF1γ deficient Tregs and promoted the maintenance of Foxp3 expression in TIF1γ cKO Tregs (Fig 4g). All together, these data suggest that the acquisition of an increased proliferative capacity and promotion of Foxp3 loss, as well as the acquisition of a Th1-like phenotype on TIF1γ cKO Tregs depends on the accumulation of β-catenin and potential increased activation of its signaling pathway.

**Fig 4.**
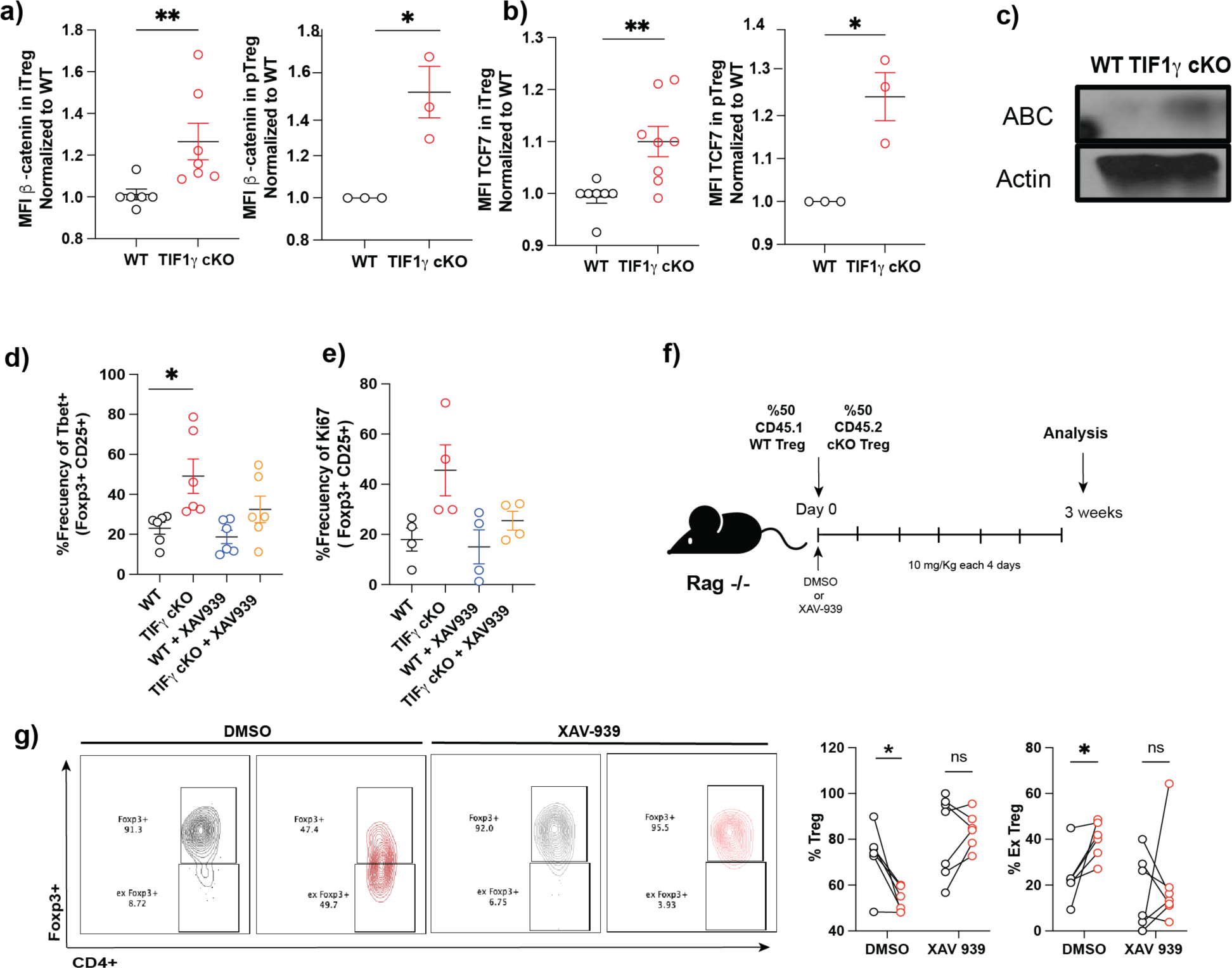
TIF1γ regulates β-catenin levels in Tregs to inhibit their proliferative capacity and acquisition of a Th1-like phenotype. a) Mean fluorescence intensity of β-catenin in iTregs and pTregs from WT or TIF1γ cKO mice. b) Mean fluorescence intensity of TCF-7 in iTregs and pTregs from WT or TIF1γ cKO mice. c) Active β-catenin (ABC) in lysates from WT or TIF1γ iTregs. d) Frequencies of Foxp3+Tbet+ in WT or TIF1γ cKO iTregs in the presence or absence of the β-catenin inhibitor XAV939. e) Frequencies of Ki67^+^ cells in WT or TIF1γ cKO iTregs in the presence or absence (DMSO as vehicle) of the β-catenin inhibitor XAV939 (11 nM). f) Experimental design of co-transferred WT (CD4^+^CD25^+^GFP^+^CD45.1^+^) and TIF1γ cKO (CD4^+^CD25^+^GFP^+^CD45.2^+^) Tregs into Rag^-/-^ hosts i.p injected every 4 with 10mg/kg of XAV939 or vehicle. g) Representative contour plots of in vivo XAV 939 treatment in co-transfer experiments. Frequencies of WT or TIF1γ cKO Tregs and exTregs in the spleen at 3 weeks post-transfer into Rag^-/-^ hosts treated with XAV939. n= 6-7, (except a and b MFI were n=3 mice per phenotype was used). Results from at least two-three independent experiments. p values: *p< 0.05, **p< 0.01

## Discussion

The TGF-β canonical pathway has been extensively shown to be a critical regulator of Treg homeostasis, differentiation and stability (3–8,12–15, 36); however, the role of the alternative TIF1γ-dependent pathway in Tregs has been largely ignored. Most of the T cell conditional deficient mice in the TGF-β signaling pathway, have shown an autoinflammatory phenotype that has been largely associated to an increased inflammatory potential of effector T cells, with minor or no apparent defect in Treg function at steady state or upon challenge (7,8, 14–15, 36–41). Our data suggest, that similar to what has been reported for Tregs deficient in TGF-β signaling pathway components; TIF1γ cKO Tregs showed no major differences compared with WT Tregs at steady state; however, unlike any other previous reports eliminating different TGF-β molecular components in Tregs, deletion of TIF1γ in Tregs severely impaired Treg function and stability upon inflammation or Treg activation.

While the role of TIF1γ in the immune system has been previously explored in HSCs, NKT cells, macrophages, Th17 cells, and more recently DCs (1, 16, 42–44); the relevance of TIF1γ for Treg function and stability has been largely ignored. Previous attempts to demonstrate a role for TIF1γ in T cell development using a CD4-Cre TIF1γ cKO mice, showed no effect on T cell differentiation in the thymus mostly focusing on CD4 and CD8 SP development, while NKT cell development was partially impaired (42). Subsequent studies in which the TGF-β signal was explored in negative selection, using TGFβRI, TGFβRII, Smad4 and TIF1γ, confirmed no apparent effect of TIF1γ ablation in T cells (45); however, none of these studies analyzed the Treg compartment, probably given the fact that these mice did not show any signs of inflammation at steady state. More recently, a new study trying to characterize the role of TIF1γ in T cell differentiation and effector subsets, confirmed no apparent phenotype at steady state in the thymus, with normal frequencies of CD4 and CD8 cells as well as no autoimmune phenotype in these mice (16), the authors did not show any quantification of Treg subsets. The same study explores the role of TIF1γ in the TGF-β dependent T cell effector subsets Th17 and Treg. The authors did not find differences in the frequencies of iTreg cells generated *in vitro*; however, Th17 cells seem to depend on TIF1γ to maintain IL-17 expression and to avoid acquisition of IL-10. Using Foxp3 Cre TIF1γ cKO mice, acquisition of IL-10 is also observed in Tregs during an experimental autoimmune encephalomyelitis model (16); although no further characterization of this phenotype was explored. This data, together with previous data on HSCs (1), centers TIF1γ as a factor able to regulate cell fate or lineage determination in immune cells; either by promoting erythrocyte differentiation in HSCs or inhibiting IL-10 expression in Th17 and Treg effector cells.

Our data further confirmed that young TIF1γ cKO mice, had no obvious phenotype in homeostasis; although it shows for the first time, that Tregs from a CD4cre TIF1γ flox/flox mice have impaired expression of CD25 and CD73, as well as a decreased activated phenotype (CD62L^-^ CD44^+^) in peripheral lymphoid organs. Interestingly, analysis of TIF1γ cKO aged mice showed signs of increased inflammation, with shortened colon length, increased weight loss, decreased activated Tregs, and increased expression of IFNγ in splenic Tregs compared to WT littermates. Expression of INFg on Tregs (Th1 like-Tregs) has been associated with an increased suppressor capacity towards Th1 cells and CTLs (24), then we decided to explore whether Tregs lacking TIF1γ expression showed any difference in their suppressor capacity. In a first set of experiments, we performed both *in vitro* suppressor assays with pTregs directly isolated from TIF1γ cKO mice, as well as an *in vivo* tumor model challenging the TIF1γ cKO mice with B16 melanoma cells. The suppressor capacity of TIF1γ cKO Tregs evaluated *in vitro,* was significantly impaired compared to WT Treg co-cultures, suggesting that TIF1γ might be regulating the suppressor function of Tregs *in vivo*. Importantly, the reduced tumor growth as well as the proper homing of Tregs to the tumor in the melanoma model and their activation status, provided additional evidence suggesting a defective suppressor function of TIF1γ deficient Tregs. However, due to the nature of the TIF1γ cKO mice (CD4 cre) used in these experiments, we cannot exclude the possibility of potential effects of lacking TIF1γ in other T cells these cells that could further contribute to the increased anti-tumor response observed in TIF1γ cKO mice.

While previous studies discarded a relevant role for TIF1γ in iTreg differentiation (16), our findings demonstrated that TIF1γ is not only important to maintain Foxp3 and prevent IFNg expression in Tregs, but it also restrains the proliferative capacity of these cells. The metabolic profile of T effectors cells has been determined in recent years (46), and is generally accepted that Treg cells are mostly dependent on oxidative phosphorylation and fatty acid oxidation, while Th1 cells have a glycolytic metabolic profile (47–48). We decided to explore if the acquisition of a Th1-like Treg phenotype observed in TIF1γ deficient Tregs, was also accompanied by changes in the metabolic profile of these cells. Strikingly, the gain of a Th1-like Treg signature in TIF1γ cKO Tregs, appears to be associated with an increased glycolytic capacity, further suggesting that TIF1γ is important for the maintenance of Treg identity both at the metabolic level and the effector level. Our data goes in agreement with recent studies relating both, the TGF-β signaling and the TIF1γ molecule in the regulation of the metabolic profile observed in human T cells and erythrocyte differentiation (49–50). It also supports previous studies where an enhanced glycolytic profile in Tregs is associated with decreased Foxp3 expression and loss of stability (48, 51–53).

Our results suggest that TIF1γ becomes relevant in Treg biology in a context-dependent manner, since a clear phenotype of its absence was only observed upon Treg activation or in the presence of an inflammatory environment. Accordingly, we believe that this is the reason why previous studies did not find any apparent phenotype at steady state in TIF1γ cKO young mice, and we only observed some signs of phenotypic differences in aged mice, where homeostatic inflammation takes place. To explore this idea and to provide definitive evidence that TIF1γ is required in a context-dependent manner, we evaluated the loss of Foxp3 expression *in vitro* by activating Tregs in the presence or absence of inflammatory cytokines. Our data confirm our hypothesis that TIF1γ cKO Tregs are especially susceptible to lose of Foxp3 expression in the presence of an inflammatory stimuli. Furthermore, we performed *in vivo* adoptive cell transfer (ACT) experiments where we explored the functional capacities and phenotypic characteristics of TIF1γ cKO Tregs. Initial experiments using the ACT colitis model showed that the suppressor capacity of TIF1γ cKO Tregs was severely impaired, since mice co-transfered with cKO Tregs showed inflammation signs and colon tissue destruction indistinguishable from the single-transfered naive control group. Remarkably, analysis of Treg populations in these experiments demonstrated an increased frequency of ex-Tregs (Foxp3^-^) and a decreased frequency Foxp3+ cells in TIF1γ cKO Tregs compared to WT controls. Additionally, the acquisition of IFNγ expression in both Tregs and ex-Tregs, was significantly increased in TIF1γ cKO donor Treg cells. We further demonstrated that the phenotype observed in TIF1γ cKO Tregs was cell intrinsic and could not be prevented by bonafide Treg cells. We also showed that the low methylation status of the CNS2 enhancer region in the Foxp3 locus, a hallmark characteristic of a stable Treg, was increased in the absence of TIF1γ, importantly this difference was acquired only after Treg differentiation because naive TIF1γ cKO cells did not have such increased methylation at this enhancer region.

TIF1γ has been involved both in TGF-β signaling but also as part of a degradation pathway for β-catenin, mostly dependent on its E3 ubiquitin ligase domain (1,17). To explore if the TGF-β signaling was responsible for the phenotype observed in TIF1γ cKO Tregs, we performed *in vitro* experiments either promoting or inhibiting the TGF-β pathway. Our data suggest that the acquisition of a Th1-like Treg phenotype in TIF1γ cKO cells was further enhanced in the absence of TGF-β, whereas the increased proliferative capacity observed in TIF1γ deficient Tregs was partially independent. Previous studies using TGFβRI, TGFβRII or TGFβ conditional deficient mice (7–8, 37), have shown an increased capacity of Treg cells to acquire a Th1-like phenotype, further supporting a critical role for TGF-β signaling in the inhibition of Th1 related genes in Tregs (7, 37). There is also previous work suggesting that Smad4 is important to promote T cell proliferation (13); here, we demonstrated that deletion of TIF1γ in T cells have the opposite effect on proliferation, supporting a competitive model for Smad4 and TIF1γ dependent signals (1), in the regulation of proliferation in T cells.

On the other hand, analysis of the β-catenin pathway in TIF1γ cKO Tregs demonstrated that in the absence of TIF1γ, Treg cells accumulate β-catenin and showed increased expression of TCF7, a downstream target of β-catenin. We also demonstrated that TIF1γ deficient Tregs have increased active β-catenin compared with wt Tregs, further suggesting that decreased degradation of β-catenin is occurring in the absence of TIF1γ. To address the relevance of increased β-catenin activation in the phenotype observed in TIF1γ cKO Tregs, we used XAV939 to activate the canonical degradation complex of β-catenin. Our data demonstrated that both acquisition of a Th1-like phenotype as well as increased Treg proliferation, was ablated when the levels of β-catenin were decreased. These results are in agreement with previous work where expression of a constitutively active form of β-catenin in Tregs, is enough to promote a wasting syndrome in mice, increased expression of IFNγ, decreased suppressor capacity, loss of Foxp3 expression and increased proliferation in Tregs (25, 31–32); then the absence of TIF1γ in Tregs, largely overlaps with the phenotype observed by a constitutively active form expressed in Tregs. To further validate that this pathway was indeed responsible for the phenotype observed in TIF1γ cKO Tregs, we repeated the co-transfer experiment of WT and cKO Tregs in Rag ^-/-^ mice, in the presence of XAV939. Our results demonstrated that inhibition of β-catenin accumulation in this system is enough to ablate the phenotype of instability in Tregs *in vivo*.

Altogether, our work demonstrates a critical role of TIF1γ for Treg stability and maintenance of suppressive functions in a context-dependent manner *in vivo.* Importantly, the fact that TIF1γ was only required to stabilize a Treg phenotype during inflammatory conditions or upon Treg activation; which are the scenarios where Treg function is critically required, further centers TIF1γ as an interesting candidate to modulate Treg function and plasticity for therapeutic purposes.

## Acknowledgements

P.L-L. is supported by UNAM (PAPIIT-DGAPA) IN220823 and Conahcyt Pronace 303027. E.C-C received a fellowship from Conacyt 255287 and PAPIIT. J.D.Z-R was supported by Conahcyt fellowship 931997. E.C-C conducted this study to fulfill the requirements of Programa de Doctorado en Ciencias Bioquímicas at Universidad Nacional Autónoma de México. We would like to thank the Animal Facility, the Molecular Biology Unit, the Microscopy unit and the Histology unit from the Instituto de Fisiología Celular for technical support. We would also like to thank technical support from Damaris Priscilla Romero Rodriguez at the Flow cytometry Unit-INER and from Jessica Prieto at the Flow Cytometry Lab-CMNSXXI.

## Author Contributions

P. L-L. designed and conceptualized the project, performed the experiments, analyzed data, provided resources and wrote the manuscript. E.C-C conceptualized the project, performed the experiments, analyzed data, and wrote the manuscript. J.D.Z-R and D.P-V discussed the data and performed experiments. I.I.L-L and M.G.G-P analyzed, discussed the data and reviewed the paper. G.T-U and F.R-T performed, analyzed and discussed the bisulfite sequencing experiments. J.L.R-B. technically assisted the project, helped with administrative duties and discussed the results. F.R-S, J.C.C-A and J.H-M discussed the results and provided technical support. I.M-S and L.N-L helped with the Seahorse experiments. A.C-M and P.N provided technical support and reagents for the β-catenin analysis. V. K. kindly donated the Trim33 Flox/Flox mice (18). R.F. provided initial resources and discussed the project.

## Methodology

### Animals

All animals were on a C57BL/6 background. TIF1γ cKO mice were generated by crossing the Trim33 Flox mice (Kindly donated by Dr V Kaartinen (18)) with the B6.Cg-Tg(Cd4-cre)1Cwi/BfluJ (19) (Cd4Cre, Jackson Labs). The mouse strains B6.Cg-Foxp3tm2Tch/J GFP, B6(129S4)Gt(ROSA)26Sortm1.1(CAG-tdTomato/GCaMP6f)Mdcah/J, B6.SJL-PtprcaPepcb/BoyJ and B6.129(Cg)-Foxp3 tm4(YFP/icre)Ayr/J were purchased from Jackson Labs. All experimental mice were generated and bred at the core animal facility of the Instituto de Fisiologia Celular (IFC). For the colitis model B6.129S7-Rag1tm1Mom/J (Rag1-/-) (Jackson labs) were used. Animals were bred and maintained under specific pathogen-free conditions in accordance with IACUC- IFC committee regulations (PLL-220-23) and CICUAL-INCMNSZ (IRE-2092). All studies were performed on gender-matched mice 8-14 weeks of age. Aged mice were housed and after reaching 40 weeks old the animals were euthanized for analysis. Both male and female mice were included in the study and we did not observe sex-dependent effects. All mice analyzed were age and litter-matched unless otherwise specified.

### Suppression assays

Suppression assays were performed by coculturing 1 × 10^5^ Cell Trace Violet (CTV) labelled and previously FACS sorted CD45.1^+^ CD4^+^ CD25^-^ CD62L^+^ CD44^-^ naive cells with varying ratios of FACS sorted CD45.2^+^ CD4^+^ CD25^+^ Foxp3^+^(GFP) Treg isolated from either FoxP3-GFPWT or FoxP3GFP-cKO TIF1γ mice in the presence of 5 × 10^5^ MMC treated splenocytes (Splenocytes were incubated (37°C, 5% CO2) for 30 minutes with MMC (40 μg/mL) in RPMI 1640 culture medium containing 10% fetal bovine serum and penicillin/streptomycin, were then washed 3 times and resuspended in fresh RMPI 1640 mediumand 0.5ug/ml of soluble anti-CD3. Dilution of CTV was measured by flow cytometry after 5 days of coculture.

### iTreg and pTreg cultures

For iTreg cultures, naive CD4+ T cells from the spleen, axillary and inguinal lymph nodes of WT and TIF1γ cKO mice were enriched using a EasySep mouse naive CD4 T cell isolation kit (STEMCELL). Then, naive CD4+ T cells (CD25^−^CD62L^+^CD44^−^GFP^-^) were purified by flow cytometry sorting and cultured for 5 days in Clicks medium (Sigma-Aldrich) supplemented with 10% FCS, L-glutamine (2 mM), penicillin (100 U/ml), streptomycin (100 μg/ml), and β-mercaptoethanol (40 nM) in the presence of plate-bound anti-CD3 (clone 2C11, at 5 ug/ml) and anti-CD28 (clone 37.51, at 2 ug/ml), 1 ng/ml TGF-β and IL-2 (100 u/ml). For isolation and activation of pTregs, total CD4^+^ T cells from the spleen, axillary and inguinal lymph nodes of WT and TIF1γ cKO mice were enriched using an EasySep Mouse CD4 T cell isolation kit (STEMCELL) followed by a purification by sort of CD4^+^CD25^+^Foxp3^+^(GFP) cells. Purified cells were cultured for 5 days in Clicks medium (Sigma-Aldrich) supplemented with 10% FCS, L-glutamine (2 mM), penicillin (100 U/ml), streptomycin (100 μg/ml), and β-mercaptoethanol (40 nM) in the presence of plate-bound anti-CD3 (5 ug/ml) and anti-CD28 (2 ug/ml) and IL-2 (200 u/ml). For a set of activation experiments pTreg were additionally exposed to IFNγ (50 ng/ml), TNFα (50 ng/ml), IL-4 (20 ng/ml) or IL-6 (10 ng/ml). For neutralization experiments anti-TGF-β (10 ug/ml) were used. TGF-β (1 ng/ml) was used for some in vitro experiments together with plate bound anti-CD3 and anti-CD28. Experiments with XAV-939 (11 nM) were performed in iTreg cultures as described above for 5 days.

### T-cell transfer experiments

For the T-cell transfer colitis model, 4x105 sorted naive CD4 T cells CD4^+^CD25^-^CD62L^+^CD44^-^ CD45.1^+^ were injected i.v. into Rag 1^-/-^ recipients with or without 1 x 105 sorted pTregs (CD4^+^CD25^+^GFP^+^CD45.2^+^) from WT or TIF1γ cKO mice. Mice were killed at indicated time points or killed when weight loss approached 20% of the original body weight at the start of the experiment. In co-transfer experiments, 5-10 x 104 sorted WT pTregs (CD4^+^CD25^+^YFP^+^CD45.1^+^) or TIF1γ cKO pTregs (CD4^+^CD25^+^YFP^+^CD45.2^+^) from CD4 Cre and fate Foxp3Cre mice were mixed and injected i.v. into Rag 1^-/-^ hosts; in some experiments, i.p administration of XAV 939 (10 mg/kg) was performed every 4 days until analysis. For single transfer experiments 5-10 x 10^4^ sorted fate WT pTregs (CD4^+^CD25^+^YFP^+^tdtTomato^+^) or fate TIF1γ cKO pTregs (CD4^+^CD25^+^YFP^+^tdtTomato^+^) were injected i.v. into Rag 1^-/-^ hosts. For pTreg induction *in vivo*, naive CD4 T cells CD4^+^CD25^-^ CD62L^+^CD44^-^ from WT or TIF1γ cKO mice were single-transferred i.v into Rag 1^-/-^ recipients and analyzed at 3 weeks post-transfer.

### Lymphocyte Preparation

Spleen, inguinal lymph nodes, axillary lymph nodes, mesenteric lymph nodes, and thymus were removed from euthanize mice, placed into RPMI media supplemented with 10% fetal bovine serum (FBS), and passed through a cell strainer (70 μm). For isolation of TILs, cells were collected after 1 hr digestion in RPMI media supplemented with 5% FCS, 1 mg/ml Collagenase D (Roche), and 100ug/ml DNase I (Invitrogen) at 37°C. Homogenates were passed through a cell strainer, and infiltrates were separated with a 27.5% Optiprep gradient (Axis-Shield) by centrifugation at 2,750 rpm for 20 min. Cells were removed from the interface, and ammonium-chloride-potassium lysing buffer was used. Cells were stained for surface markers and/or intracellular antigens. All flow cytometry data were acquired on a Attune Next (Thermo Fisher Scientific) or Aurora (Cytek) and analyzed by FlowJo software (v10.8.1, Treestar).

### Flow cytometry staining and cell sorting

The full list of antibodies used in this study are shown in Extended table I. Live cells were discriminated using Live/Dead Fixable Aqua Dead Cell Stain Kit (Biolegend) in PBS according to the manufacturer’s protocol. Surface staining was performed for 30 min at 4 °C in FACS Buffer (PBS + FBS 2%). Intracellular staining for transcription factors was conducted using the Foxp3 Transcription Factor Buffer Set (Life Technologies) according to the manufacturer’s protocol. Cell sorting was performed on a FACS Aria II and FACS Melody (BD Biosciences). Enriched CD4+ T cells were sorted to over 95% purity on FACS Aria II and FACS Melody (BD Biosciences). Sort was performed with the following gates after excluding death cells CD4^+^CD25^+^YFP/GFP^+^ and CD4^+^ CD62L^+^ CD44^-^ GFP^−^ for naive T cells.

### Intracellular cytokine staining

For cytokine production by Treg cells, lymphocytes were stimulated with 50 ng/ml of phorbol 12-myristate 13-acetate (PMA) and 1000 ng/ ml of ionomycin and GolgiStop (Bioscience) in supplemented RPMI 1640 and incubated for 4 hours (37°C 5% CO2). After stimulation, cells were stained for viability staining and surface staining. Intracellular staining for cytokines was conducted in Perm/Wash buffer after fixation and permeabilization in Fix/ Perm buffer (BD Biosciences) according to manufacturer instructions.

For flow cytometric analysis the following gating strategies were used to identify cell populations: for iTreg or pTreg cultures single, live, lymphocytes (based on FSC, SSC and live/dead fixable dye Aqua stain), CD4, CD25 and Foxp3 (GFP or YFP) markers were used. For *in vitro* Treg suppression assays CD45.1 and CD45.2 was added to the surface staining mix. For B16 tumor models the following gating strategies were used to identify cell populations by FACS: for lymphoid organs: single, live, lymphocytes (based on FSC, SSC and live/dead fixable dye Aqua stain), CD8, CD4, CD3, Foxp3 (GFP), CD25. For GFP^+^ cells additional surface markers were analyzed CD44, CD62L, CD73, CD39 and PD-1. For TILs: single, live, lymphocytes (based on FSC, SSC and live/dead fixable dye Aqua stain), CD45.2, CD8, CD4, CD3, Foxp3 (GFP) and CD25 were used. For T cell transfer experiments, the following gating strategies were used to identify cell populations by FACS: T cells: single, live, lymphocytes (based on FSC, SSC and live/dead fixable dye Aqua stain), CD45.1/CD45.2, TCRβ, CD4, CD3 and Foxp3. Ki67, Tbet, β-catenin and IFNγ was also analyzed in Fox3^+^ and Foxp3^-^ populations. For some experiments tdtTomato^+^ was also considered as a first marker for gating strategy before Foxp3^+^ (YFP) cells.

### Extracellular flux analyses

iTregs from 5 day cultures were sorted (CD4^+^Foxp3^+^CD25^+^) and OCR and ECAR values were measured using a 96-well XF Extracellular flux analyzer, following manufacturer instructions (Agilent).

### Proliferation analysis

Sorted pTreg or Naive cells were stained with Tag-it Violet reagent according to the manufacturer instructions (Biolegend) after 5 days culture. pTregs of iTregs were harvested stained for surface markers and viability dye Zombie . Proliferation analysis was performed following the dilution of Tag-it Violet by flow cytometry. Proliferation index was calculated using the FlowJo software.

### Apoptosis analysis

*In vitro* differentiated iTregs were harvested, stained for surface markers and viability dye Zombie. Stained cells were incubated with Annexin V (Biolegend) according to the manufacturer instructions.

### Sodium bisulfite DNA conversion and sequencing

Cells were lysed in Lysis Buffer (100 mM Tris-HCl pH 8.0, 200 mM NaCl, 0.2% SDS, 5 mM EDTA) in presence of proteinase K (incubation at 55 °C for five hours), and genomic DNA was extracted with 1:1 Phenol:Chloroform:Isoamyl Alcohol (Invitrogen) and precipitated with 2-propanol. Bisulfite conversion of DNA was conducted as described in Clark et al. (54) with some modifications. Briefly, 2 µg of DNA were denatured at 95°C for 5 min, chilled on ice, and incubated with 0.3 M NaOH at 37 °C for 5 min. Freshly-prepared solutions of sodium bisulfite (Sigma), adjusted to pH 5.0 with NaOH, and hydroquinone (Sigma) were added at final concentrations of 1.7 M and 0.5 mM, respectively. DNA solutions were mixed and incubated at 55 °C for 12 h (protected from light). Non-reacting bisulfite was removed by column purification (Wizard DNA Clean-Up System; Promega). Purified DNA samples were de-sulfonated with NaOH at a final concentration of 0.3 M, incubated at 37 °C for 15 min followed by precipitation with ethanol and ammonium acetate pH 7.0 at a final concentration of 0.4 M. DNA regions of interest were amplified by nested PCR with HotStartTaq DNA Polymerase (QIAGEN) with primers designed using the MethPrimer software (55) and listed in extended Table II. PCR products were recovered after agarose gel electrophoresis with QIAEX II Gel Extraction Kit (QIAGEN), then cloned into the pGEM-T Easy system (Promega), purified, and sequenced using the T7 sequence primer.

### ATAC-seq data retrieval and processing

Publicly available ATAC-seq were obtained from the Gene Expression Omnibus (GEO) database. Raw data from different Treg cells phenotypes isolated from mouse thymus were obtained under the following accession numbers: Resting Treg (GSM4677045-GSM4677048), Activated Treg (GSM4677049 - GSM4677052), and Treg (GSM3428984-GSM3428987) (22, 30). Raw files were imported into Galaxy Europe (usegalaxy.eu) for fastq files extracting, mapping, and visualizing. Reads were trimmed using Trim Galore! (56) with default parameters. Paired-end reads were mapped against the Mus musculus genome (mm10) using Bowtie2 (57) with default parameters. Aligned reads were filtered using BAM filter (58) based on criteria, keeping only mapped properly paired reads, and removing: secondary alignment reads, reads that do not pass the quality control, reads that are marked as PCR duplicates, and reads found in the blacklist (mm10-blacklist.v2.bed; 59). Bam files from experiments with replicates were merged using MergeSamFiles (Pikard) with default parameters. Average read coverage from merged bam files was calculated with multiBAMSummary (60) with parameters: sample order does not matter, computation mode in bins of 10000 bp and saving the scaling factors between files. Merged bam files were used to create signal tracks using bamCoverage (60) in bins of 50 bases scaling with the scaling factor computed in the previous step. Signal tracks were visualized using the UCSC Genome Browser.

### Western blot analysis

Sorted 1 x 10^6^ iTregs from WT or TIF1γ cKO were collected and homogenized in RIPA buffer containing 150 mM NaCl, 1% NP-40, 0.5% deoxycholic acid, 0.1% SDS, 50 mM Tris, at a pH of 8.0. The protein concentration was determined using the BCA Protein Assay (Pierce, Thermo Fisher Scientific). Subsequently, 10 µg of protein were loaded onto a denaturing SDS-PAGE for analysis. For active beta catenin (ABC) detection the antibody from Cell signalling: Non-phospho (Active) β-Catenin (Ser33/37/Thr41) (D13A1) Rabbit mAb #8814 was used according to manufacturer instructions. For actin detection, a Mouse anti-actin antibody was used, kindly donated by Dr. Manuel Hernández; Department of Cell Biology, CINVESTAV.

### Tissue Histology

Colon tissue from colitis experiments were harvested, cleaned and fixed in 4% PFA overnight. The tissue was embedded in paraffin and sections were stained with Haematoxilin and eosin.

### Statistical analysis

Details on statistics used can be found in figure legends. Data were subjected to the 905 Kolmogorov-Smirnov test to assess normality of samples. Statistical differences were calculated by using unpaired two-tailed Student’s t-test, paired two-tailed Student’s t-test and/or one-way ANOVA with Tukey post-test. All experiments were analyzed using Prism 10 (GraphPad Software).

Graphical data was shown as mean values with error bars indicating the Standard Deviation. p values of ≤ 0.05 (*), ≤ 0.01 (**), ≤ 0.001 (***) or ≤ 0.0001 (****) indicated significant differences between groups.

