## Extended figures and tables for "TIF1γ regulates stability of T regulatory cells during inflammation"

**Extended data figure legends**

**Extended data Figure 1. TIF1γ deficient T cells have an impaired iTreg differentiation capacity and proliferate in the presence of inflammatory stimuli.** a) Frequency of $CD4^+$, $CD8^+$ and $CD4^+Foxp3^+$ cells in peripheral lymph node (PLN), mesenteric lymph node (MLN) and spleen of 8-week-old WT and TIF1γ cKO mice. b) Mean fluorescence intensity of CD25 and CD73 in $CD4^+Foxp3^+$ cells from PLN, MLN and spleen of 8-week-old WT and TIF1γ cKO mice (data normalized to WT). c) Frequencies of $Foxp3^+IFN\gamma^+$ cells in the spleen of 40-week-old WT and TIF1γ cKO mice. d) Frequencies and absolute numbers of iTregs generated *in vitro* at day 5 from naive WT or TIF1γ cKO cells. e) Frequencies of $Foxp3^+IFN\gamma^+$ and $Foxp3^+Tbet^+$ cells in iTreg cultures at day 5 from WT or TIF1γ cKO mice. f) Proliferation index in iTreg cultures from WT or TIF1γ cKO mice. g) Apoptosis analysis in iTreg cultures from WT or TIF1γ cKO mice, percentages of live ($AnexinV^-Zombie^-$), apoptotic ($AnexinV^+Zombie^-$) and death ($AnexinV^+Zombie^+$) cells in cultures. h) Frequencies of Foxp3+CD25+ and mean fluorescence intensity of Foxp3 in pTregs from WT or TIF1γ cKO mice stimulated *ex vivo* with anti CD3, anti-CD28, IL-2 and different inflammatory cytokines at day 5. i) Representative histograms of CTV dilution (proliferation) in pTReg cultures from WT or TIF1γ cKO mice stimulated *ex vivo* with anti CD3, anti-CD28, IL-2 and different inflammatory cytokines at day 5. j) CD8 to Treg ratio in tumor infiltrating lymphocytes (TILs) from WT or TIF1γ cKO mice at day 21 post B16 inoculation. k) Mean fluorescence intensity of CD25, CD73, CD39 and PD1 in Tregs from spleen, draining lymph nodes (DLN), non-draining lymph nodes (NDLN) and TILS of WT or TIF1γ cKO mice at day 21 post B16 inoculation. n= 4-13 from at least three independent experiments. p values: *p< 0.05, **p< 0.005, ****p<0.0001

**Extended data figure 2. TIF1γ deficient Tregs expand *in vivo* and lose Foxp3 expression upon transfer into Rag-/- hosts.** a) Normalized mean fluorescence intensity of Foxp3 expression in splenic Tregs from WT or TIF1γ cKO mice co-transfered with naive T cells into $Rag^{-/-}$ hosts at day 52 post transfer. b) Body weight change of $Rag^{-/-}$ mice single transferred with WT or TIF1γ cKO fate Tregs. c) Fold increase in Treg and exTreg numbers from spleen of $Rag^{-/-}$ mice single transferred with WT or TIF1γ cKO fate Tregs at day 52 post transfer. d) Experimental design for pTreg induction *in vivo* in $Rag^{-/-}$ hosts transferred with naive fate WT or TIF1γ cKO cells. e) Representative contour plot of Tregs and exTregs in spleen of $Rag^{-/-}$ mice transferred with naive WT or TIF1γ deficient T cells. Frequencies of Tregs and exTregs in spleen and MLN of $Rag^{-/-}$ mice transferred with naive WT or TIF1γ deficient T cells. f) Frequencies of fate WT or TIF1γ cKO Tregs and exTregs in the MLN at 3 weeks post-transfer into $Rag^{-/-}$ hosts. g) Frequencies and absolute numbers of WT or TIF1γ cKO (CD4 Cre) Tregs and exTregs in the spleen at 3 weeks post-transfer into $Rag^{-/-}$ hosts. h) Frequencies and absolute numbers of WT or TIF1γ cKO (CD4 Cre) Tregs and exTregs in the MLN at 3 weeks post-transfer into $Rag^{-/-}$ hosts. n= 10-11 (except b, c and e were n=3 mice per phenotype was used) from at least two independent experiments. p values: * p<0.05, ** p<0.01, *** p<0.001

**Extended data figure 3. TIF1γ deficient Tregs have an increased methylation profile at the CNS2 enhancer region of the Foxp3 locus.** a) Chromatin accessibility profiles, retrieved from public ATAC-seq data (22,30), of resting Tregs, activated Tregs and thymic Tregs (nTregs). DNA sequence conservation (by PhyloP) profile with the Promoter and CNS2 regions (black boxes) where CpG methylation was analyzed. b) Global CpG methylation (%CpG) of six individual CpG dinucleotides found at the promoter region of T naive WT and TIF1γ cKO cells. c) Global CpG methylation (%CpG) of ten individual CpG dinucleotides found at the CNS2 regulatory element of T naive WT and TIF1γ cKO cells. d) Global CpG methylation (%CpG) of six individual CpG dinucleotides found at the promoter region of WT and TIF1γ cKO iTregs. e) Global CpG methylation (%CpG) of ten individual CpG dinucleotides found at the CNS2 regulatory element of WT and TIF1γ cKO iTregs. f) Global CpG methylation (%CpG) of six individual CpG dinucleotides found at the promoter region of WT and TIF1γ cKO pTregs. e) Global CpG methylation (%CpG) of ten individual CpG dinucleotides found at the CNS2 regulatory element of WT and TIF1γ cKO pTregs. The average methylation found at each CpG position from two different DNA samples (R1 and R2) is presented and compared between WT and cKO cells (w and k, respectively). Comparisons were made by Wilcoxon test in GraphPad Prism 9.5.1 and the corresponding p values are shown.

### Extended 1

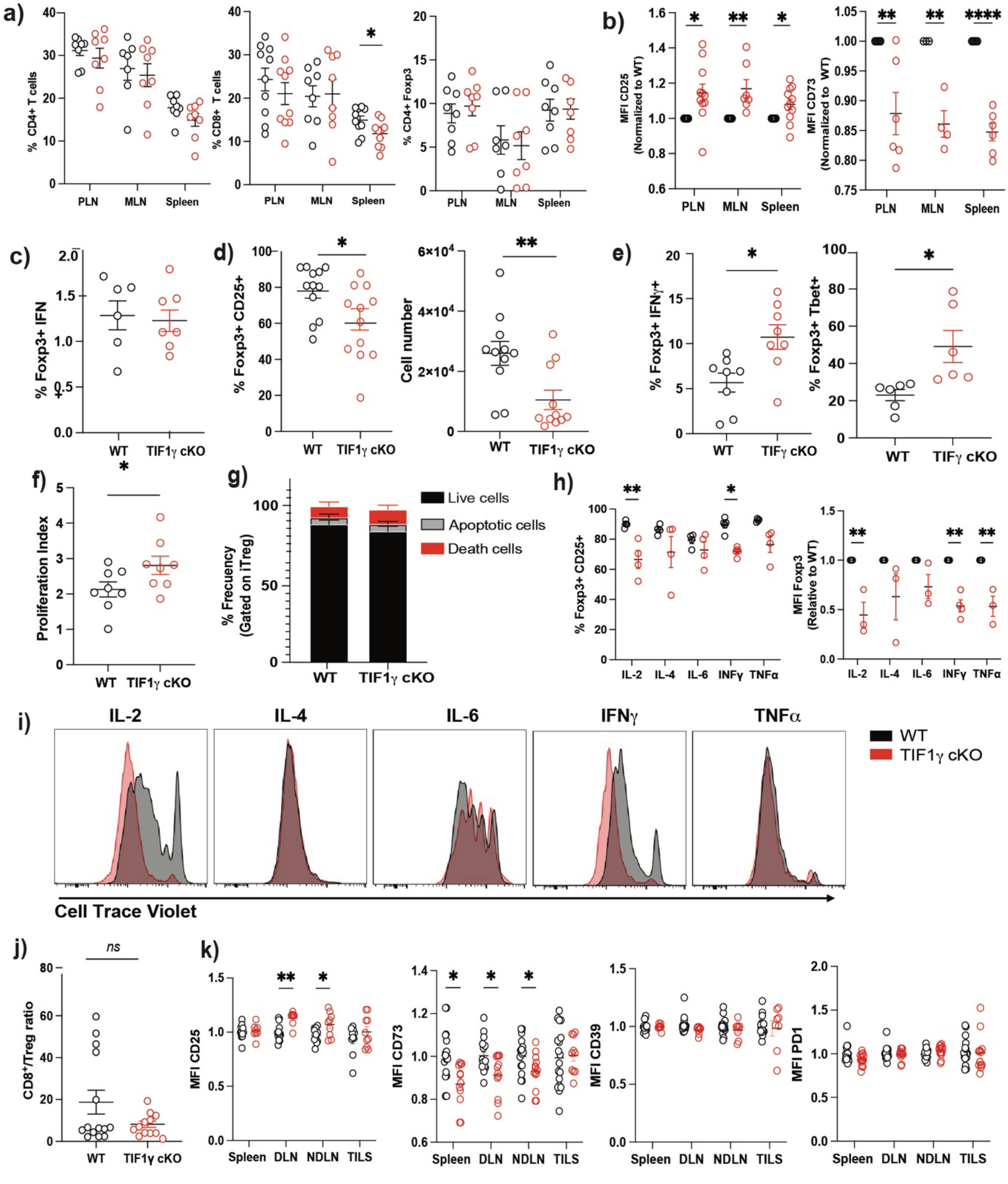

a)
% CD4+ T cells
% CD8+ T cells
% CD4+ Foxp3
PLN
MLN
Spleen
b)
MFI CD25 (Normalized to WT)
MFI CD73 (Normalized to WT)
c)
% Foxp3+ IFN
WT
TIF1γ cKO
d)
% Foxp3+ CD25+
Cell number
e)
% Foxp3+ IFNγ+
% Foxp3+ Tbet+
TIFγ cKO
f)
Proliferation Index
g)
% Frecuency (Gated on iTreg)
Live cells
Apoptotic cells
Death cells
h)
% Foxp3+ CD25+
MFI Foxp3 (Relative to WT)
IL-2
IL-4
IL-6
INFγ
TNFα
i)
IFNγ
WT
TIF1γ cKO
Cell Trace Violet
j)
ns
CD8+/Treg ratio
k)
MFI CD25
MFI CD73
MFI CD39
MFI PD1
DLN
NDLN
TILS

### Extended 2

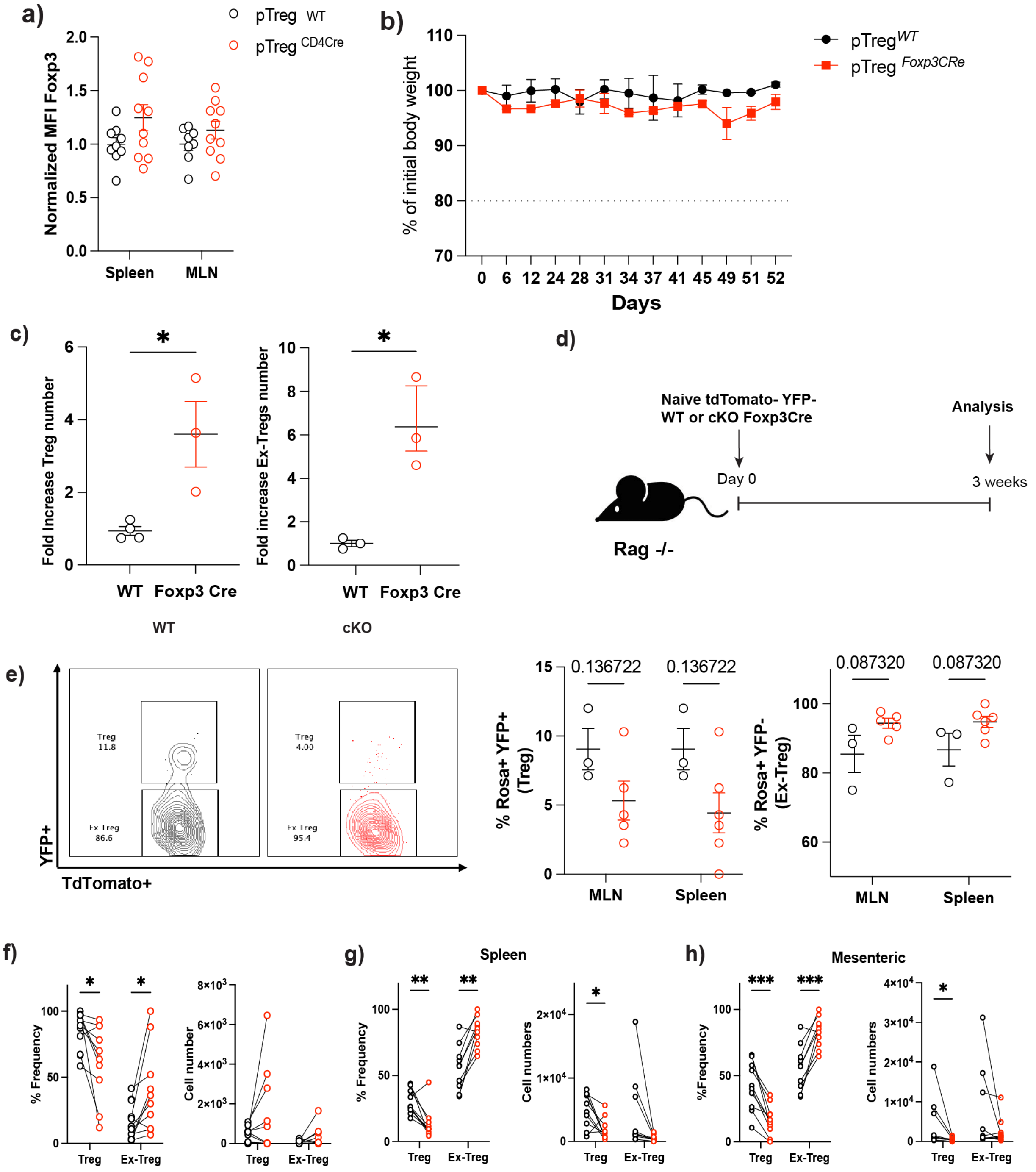

a)
pTreg WT
pTreg CD4Cre
Normalized MFI Foxp3
Spleen
MLN
b)
pTreg WT
pTreg Foxp3CRe
% of initial body weight
Days
c)
Fold Increase Treg number
Fold increase Ex-Tregs number
WT
Foxp3 Cre
WT
cKO
d)
Naive tdTomato- YFP-
WT or cKO Foxp3Cre
Analysis
Day 0
3 weeks
Rag -/-
e)
Treg
11.8
Ex Treg
86.6
Treg
4.00
Ex Treg
95.4
YFP+
TdTomato+
0.136722
% Rosa+ YFP+ (Treg)
MLN
Spleen
0.087320
% Rosa+ YFP- (Ex-Treg)
f)
% Frequency
Cell number
Treg
Ex-Treg
g)
Spleen
Cell numbers
h)
Mesenteric
%Frequency

### Extended 3

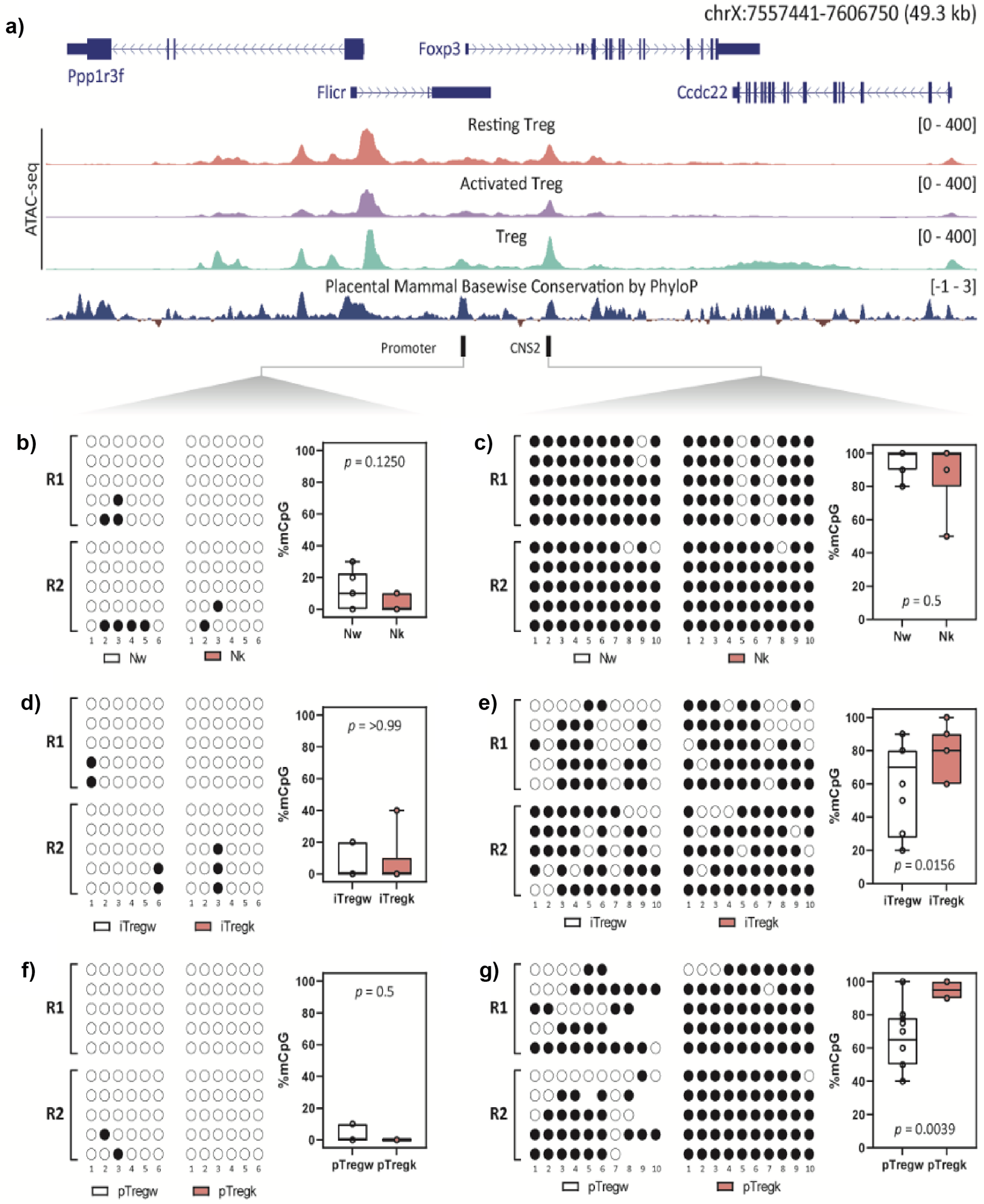
a)
chrX:7557441-7606750 (49.3 kb)
Ppp1r3f
Foxp3
Flicr
Ccdc22
ATAC-seq
Resting Treg
[0 - 400]
Activated Treg
[0 - 400]
Treg
[0 - 400]
Placental Mammal Basewise Conservation by PhyloP
[-1 - 3]
Promoter
CNS2
b)
R1
R2
%mCpG
p = 0.1250
Nw
Nk
c)
p = 0.5
Nw
Nk
d)
p = >0.99
iTregw
iTregk
e)
p = 0.0156
iTregw
iTregk
f)
p = 0.5
pTregw
pTregk
g)
p = 0.0039
pTregw
pTregk

Table I

| Anti-Mouse | Flurorofore | Catalog number | Clone |
|---|---|---|---|
| CD4 | APC | 20-0042-u100 | rm4-5 |
| cd39 | APC | 143809 | duha59 |
| Foxp3 | APC | 17-5773-82 | FJK-16s |
| CD45.1 | APC | 110714 | A20 |
| Foxp3 | AF700 | 126422 | MF-14 |
| CD4 | AF700 | 100430 | GK1.5 |
| CD45.2 | AF700 | 109822 | 104 |
| CD62L | APC/Cyanine7 | 104428 | MEL-14 |
| CD4 | APC/Cyanine7 | 100414 | GK1.5 |
| CD4 | PE | 50-0041-u100 | biolegend |
| CD73 | PE | 127206 | TY/11.8 |
| CD25 | PE | 102008 | PC61 |
| IFNg | PE | 50-7311-U025 | XMG1.2 |
| TCF1 | PE | 564217 | S33-966 |
| beta catenin 1 | PE | 862604 | 15B8 |
| CD45.2 | PE/Dazzle | 109846 | 104 |
| CD4 | PE/Dazzle | 100456 | GK1.5 |
| CD4 | PE/Cyanine5 | 100410 | GK1.5 |
| IFNg | PE/Cyanine5 | 55-7311-U100 | XMG1.2 |
| Foxp3 | PE/Cyanine5 | 55-5773-U025 | 3G3 |
| CD4 | PE/Cyanine7 | 100422 | GK1.5 |
| TCR-B | PE/Cyanine7 | 109222 | H57-597 |
| CD44 | PE/Cyanine7 | 103030 | 1M7 |
| T-BET | PE/Cyanine7 | 25-5825-82 | 4B10 |
| PD1 | PE/Cyanine7 | 135216 | 29F.1A12 |
| CD4 | Fitc | 100405 | GK1.5 |
| CD3 | Fitc | 100203 | 17A2 |
| CD45.2 | PercP | 109826 | 104 |
| CD3e | PercP/Cyanine5.5 | 65-0031-U025 | 500A2 |
| CD4 | Percp/c5 | 100434 | GK1.5 |
| CD25 | Percp/c5 | 102030 | PC61 |

| | | | |
|---|---|---|---|
| CD4 | PB | 100428 | GK1.5 |
| CD8a | PB | 100725 | GK1.5 |
| CD73 | PB | 127212 | TY/11.8 |
| CD45.1 | PB | 110722 | A20 |
| Ki67 | PB | 652422 | 16A8 |
| CD4 BV421 | BV421 | 100437 | GK1.5 |
| Annexin V | APC | 540941 | |
| Cell CellTrace™ Violet | | C34557 | |
| Zombie Aqua Fixable Viability Kit | | 423102 | |

Table II . List of oligonucleotides used in this study.

| Primer name | Sequence |
|---|---|
| Bis_Prom_F | TATATTTTTAGATGATTTGTAAAGGGTAAA |
| Bis_Prom_R | AAAAAACTACCACATTATCAAAAACAACT |
| Bis_Prom_F2 | AGATGATTTGTAAAGGGTAAAGAAAA |
| Bis_Prom_F2 | CCACATTATCAAAAACAACTTTACTT |
| Bis_CNS2_F | TTTTGGGTTTTTTTGGTATTTAAGA |
| Bis_CNS2_R | AACCAACCAACTTCCTACACTATCTAT |
| Bis_CNS2_F2 | TGGGTTTTTTTGGTATTTAAGAAAG |
| Bis_CNS2_R2 | TTAACCAAATTTTTCTACCATTAAC |
